# Glossiness perception and its pupillary response

**DOI:** 10.1101/2022.04.13.488254

**Authors:** Hideki Tamura, Shigeki Nakauchi, Tetsuto Minami

## Abstract

Recent studies have discovered that pupillary response changes depend on cognitive factors, such as subjective brightness caused by optical illusions and luminance. However, it remains unclear how the cognitive factor derived from the glossiness perception of object surfaces affects pupillary response. We investigated the relationship between glossiness perception and pupillary response through a gloss rating experiment that includes recording pupil diameter. For the stimuli, we prepared general object images (original) and randomized images (shuffled) that comprised of the same images with randomized small square regions. The image features were then controlled by matching the luminance histogram. The observers were asked to rate the perceived glossiness of the stimuli presented for 3,000 ms and changes in their pupil diameter were recorded. Consequently, if glossiness of the original images were rated as high, those of the shuffled were rated as low, and vice versa. High-gloss images constricted the pupil size more than the low-gloss ones near the pupillary light reflex. By contrast, the shuffled images dilated the pupil size more than the original image at a relatively later stage. These results suggest that local features comprising specular highlights involve the cognitive factor for pupil constriction, and this process is faster than pupil dilation derived from the inhibition of object recognition.

## Introduction

Glossiness perception is essential for estimating the surface properties of objects and plays an important role in the human visual system (Adelson, 2001; Anderson, 2011; Chadwick & Kentridge, 2015; Fleming, 2014, 2017; Komatsu & Goda, 2018). Glossy objects contain specular highlights on their surfaces (Beck & Prazdny, 1981; Fleming et al., 2003; Shinya & Nishida, 1998), and these brighter regions summarize simple features based on the luminance histogram, working as a cue of perceived glossiness (Motoyoshi et al., 2007; Nishida, 2019; Sawayama & Nishida, 2018; Sharan, Li, et al., 2008; Wiebel et al., 2015). In contrast, several studies have noted that the three-dimensional structure of an object is essential for glossiness perception (Anderson & Kim, 2009; Kim & Anderson, 2010; P. J. Marlow et al., 2012). Consequently, opinions vary on what cues the visual system uses to estimate the surface glossiness. Furthermore, the perceived glossiness is affected by components other than bright specular reflection, such as the dark region of an object surface caused by specular reflection (Kim et al., 2012; Kiyokawa et al., 2019) and image spatial frequency components (Kiyokawa et al., 2021). Thus, a simple change in luminance on the retina causes mid-level visual processing (Fleming, 2014) for glossiness perception in the visual system.

These physical and image-based characteristics of glossy objects trigger glossiness perception, which has been elucidated through neurophysiological findings. Human behavior and eye movements were associated with glossy objects (Phillips et al., 2010; Qi et al., 2018; Sharan, Rosenholtz, et al., 2008; Toscani et al., 2019), suggesting that the visual system efficiently obtains the source of information from the external world at the input stage. It is subsequently represented in the inferior temporal cortex through the ventral stream, which has been reported using functional MRI and physiological techniques in humans (Sun et al., 2015; Wada et al., 2014) and macaques (Baba et al., 2021; Komatsu et al., 2021; Nishio et al., 2012, 2014; Okazawa et al., 2012). However, the temporal dynamics linking gloss perception processing — how networks share roles in the visual system — is still unclear.

To understand this, we focused on recording the pupillary response to elicit cognitive processing affected by glossy objects. Pupillary responses reflect not only physical factors derived from luminance, but also cognitive factors such as emotional (Bradley et al., 2008; Kuraguchi & Kanari, 2020, 2021; Laeng et al., 2013) and attractive (Liao et al., 2020) stimuli. These reactions are controlled by the locus coeruleus, which affects the level of arousal (Aston-Jones & Cohen, 2005; Benarroch, 2009). In a study relevant to the relationship between pupillary response and surface properties, Tanaka et al. reported that pupil size constricts when observers carefully gaze at the material object surface (e.g., marble stone) to estimate its psychological features (Tanaka et al., 2017). Although they reported pupil responses related to physical materials and properties, little is known about whether this physiological response is derived from a physical material (e.g., stone, wood, and metal) or its surface properties (e.g., glossiness, heaviness, and colorfulness). Therefore, we focused on glossiness perception, a specific surface property, as the first step in understanding the association between pupil responses and surface properties.

Similar to arousal activities leading to pupil dilation, recent studies have revealed that cognitive factors constrict the pupil. For example, illusory brightness caused by the self-luminosity of the glare illusion causes pupil constriction as a cognitive factor (Kinzuka et al., 2021; Suzuki, Minami, & Nakauchi, 2019; Suzuki, Minami, Laeng, et al., 2019). These studies allowed us to hypothesize that changes in pupil responses are induced by glossiness perception. Specifically, specular highlight components structured by brighter regions in an image would constrict the pupil diameters. We expected that this characteristic phenomenon would occur due to one of the cognitive factors wherein we perceive an object to be glossy because the pupil diameter responds to both subjective and perceptual brightness (Laeng et al., 2018; Laeng & Endestad, 2012; Suzuki, Minami, & Nakauchi, 2019; Suzuki, Minami, Laeng, et al., 2019; Zavagno et al., 2017). If we can understand how the pupil reacts to glossy objects, the outcomes can be used to develop a new tool to estimate perceived glossiness or visual material perception without any explicit behavioral responses from human observers.

This study aimed to understand how pupil responses reflect perceived glossiness, which are modulated by image structures. Based on the above, we hypothesized that an image with highly perceived glossiness causes pupil constriction, even if its physical luminance is equalized. Alternatively, there could be no significant difference because the effectiveness of a cognitive factor influenced by perceived glossiness is smaller than that of a physical factor based on luminance. We conducted an experiment to investigate how perceived glossiness relates to pupil responses through a glossiness rating task in human psychophysics with pupillometry recording. The following stimuli for the experiment were employed: 1) general object images in our daily life and 2) shuffled pixels while maintaining the local features of the objects. Thus, we tested how perceived glossiness modulates based on the global features of the objects and how they reflect pupil responses.

## Methods

### Observers

Twenty-four naïve observers participated in the experiment. Two observers were excluded owing to eye-tracking calibration failure for one and an eyesight issue with the other. Finally, data from 22 observers were used for further analysis. Their ages ranged from 20 to 25 years (average 22.9 ± 1.2 years), and they had normal or corrected-to-normal acuity. All experimental protocols were approved by the Institutional Review Board of Toyohashi University of Technology for their use on humans in experiments, in accordance with the Declaration of Helsinki. Written informed consent for publication of their details were obtained from the study participants.

### Apparatus

The experiment was conducted in a dark booth with dim lighting (approximately 60 lx). Stimuli were displayed on a 27-inch liquid crystal display (ColorEdge 27 CS2731, EIZO) having a resolution of 1920 × 1080 pixels and refresh rate of 60 Hz (calibrated by SpyderX Elite, ImageVISION). Each observer saw the stimuli after being seated on a chair in the booth with their head secured on a chin rest to maintain a constant distance of 86 cm. Pupillary response was recorded using an eye tracker (EyeLink 1000 PLUS, SR Research) with a sampling rate of 500 Hz. A five-point calibration of the eye tracker was performed prior to each experimental session. Stimulus presentation was controlled by MATLAB using Psychtoolbox 3.0 (Brainard, 1997; Kleiner et al., 2007; Pelli, 1997).

### Stimuli

We obtained 60 images from the THINGS database (Hebart et al., 2019, 2020), including general daily-life objects. Each image was selected from a different category that was defined in the database. We scaled them down to 512 × 512 pixels, transformed each original RGB color image to grayscale, and trimmed its background while retaining the main object. Thereafter, the image features (mean, variance, skewness, kurtosis) from the pixel intensity histogram of these images were equalized by the histogram matching of the SHINE toolbox (Willenbockel et al., 2010). We defined these 60 images as “original” (see **Figure 1A)**.

**Figure 1.**
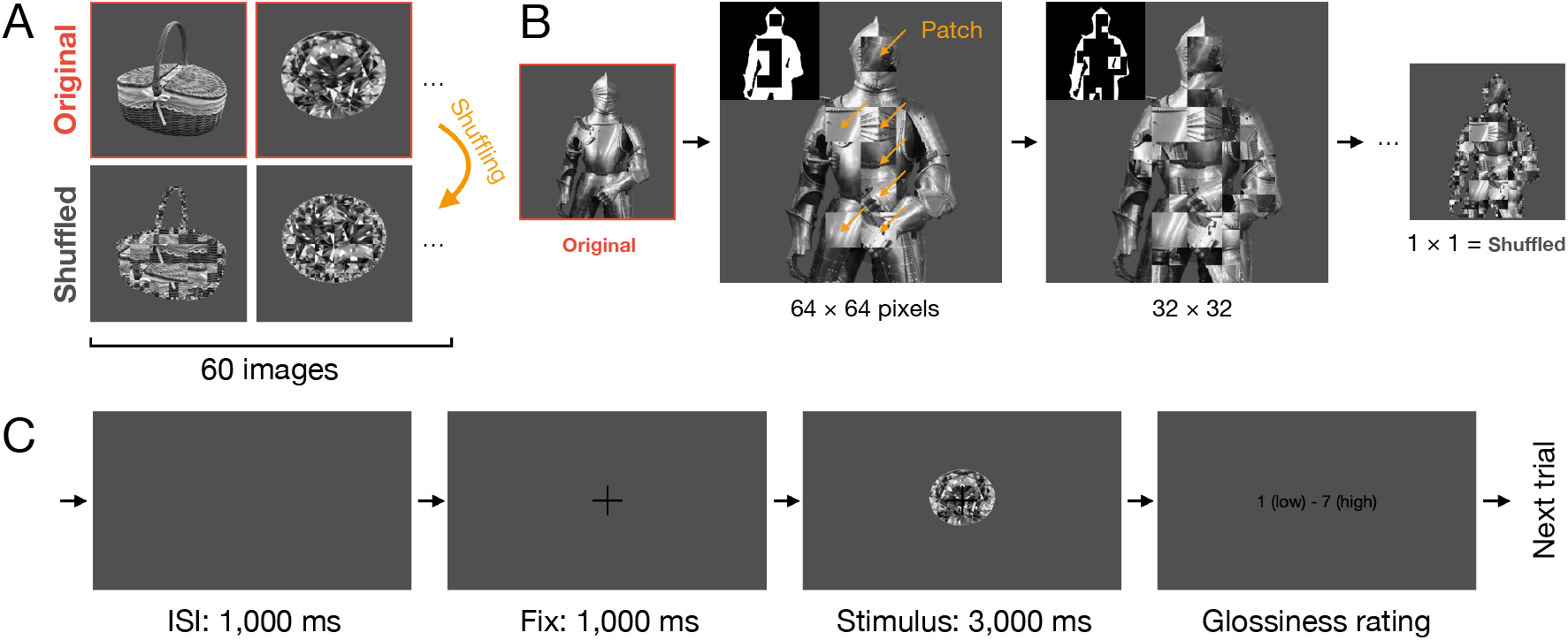
Stimuli and procedure. (A) Examples of the stimuli. Top: original; Bottom: shuffled. (B) The flow of the shuffling process. The patch size shrank in a step-by-step manner from 64 × 64 to 1 × 1 pixels. In binary images of the two middle steps, white regions indicate remaining pixels for further shuffling. (C) The sequence of one trial of the experiment. Note that the ratio of the screen to the fixation point and the stimulus in this panel differs from the actual ratio used in this study.

Additionally, a “shuffled” image was generated from each original image. **Figure 1B** illustrates the flow of this procedure. The object region of each image of 512 × 512 pixels was divided into 64 × 64 pixel patches, and the square-shaped regions were randomly shuffled. The remaining region inside the object contour was further divided into 32 × 32 pixel patches, which were then randomly shuffled. We repeated this shuffling process, starting from a patch size of 64 × 64 pixels until 1 × 1 pixel patch sizes were obtained. The output images from this manipulation maintained a local feature, whereas the global feature of the object was inhibited. Finally, we used 120 images (60 images each in original and shuffled condition) for the stimuli of the experiment.

The stimulus area was 6 × 6 degrees, and one object existed inside this area. The original and shuffled images had the same image features with the same luminance histogram because the shuffled images only scrambled the original’s pixels in different region sizes. We observed a negligible difference between the average luminance of the original and shuffled (average luminance of 60 images; original: 26.87±0.80 cd/m^2^, shuffled: 26.55±0.84 cd/m^2^), derived from scaling down the images for presentation on the experiment screen. Moreover, we verified that a local luminance change with gaze does not have any meaningful effect on pupillary response (see Discussion).

### Procedure and task

**Figure 1C** shows the trial sequence of the experiment. After a 1,000 ms inter-stimulus interval (ISI), the stimulus was presented at the screen center for 3,000 ms on a gray background (9.19 cd/m^2^) with a black fixation cross (0.6 × 0.6 degrees). Subsequently, a response-receiving screen was displayed.

The observers were asked to rate the perceived glossiness of the stimulus using a 7-point scale (1: lowest and 7: highest) with a numerical keypad. The next trial began after receiving the observer’s response. The pupil response was recorded during the experiment. Each session consisted of 120 trials (one trial × 60 objects × two image conditions), and each observer participated in two sessions separated by a break.

### Data analysis

From the observers’ responses, we computed the average glossiness ratings of all images in the original and shuffled conditions. Responses to the trials rejected through pupil preprocessing were ignored.

We analyzed pupil changes in the interval from the beginning of the fixation point to the end of the stimulus presentation. Intervals with a pupil response velocity greater than 0.011 mm/ms were considered eye blinks, including 20 ms on either side. They were then excluded from the analysis and interpolated using shape-preserving piecewise cubic spline interpolation. Furthermore, trials that included eye blinks exceeding one second, eye blinks with a ratio over 0.3 in a trial, and pupils that were not detected at the beginning or end were also excluded. An observer with a rate of rejected trials over 50% was excluded, and we used the data of the remaining 21 observers (mean and standard deviation of rejected trials was 9.2 ± 11.4%). Baseline correction was performed using the same procedure as that in Nakakoga et al. (2021). The baseline was defined as the average pupil diameter for 200 ms prior to the beginning of the stimulus presentation, and we computed each pupil’s diameter change by subtracting the baseline from the original pupil diameter. Thereafter, a moving average filter having a 10 ms window size was applied to the responses.

## Results

### Glossiness rating

**Figure 2A** shows the average glossiness ratings for each stimulus as a histogram. The ratings of the original images form two peaks and are distributed broadly (average 3.91 ± 1.33, range 1.57 to 6.14). However, the shuffled ratings show a center-peaked distribution (average 3.82 ± 0.77, range 2.20 to 6.05). These findings indicate that the perceived glossiness changes through the shuffled operation despite both having the same image features. **Figure 2B** shows that the original and shuffled conditions were significantly correlated (*r* = 0.81, *p* < 0.001), and the slope of the regression line was less than one (*y* = 0.47*x* + 1.99). These findings indicate that the glossiness ratings of the original and shuffled images are inversely related. Some of these results suggest that pixels constituting the specular highlight on the object’s highly rated gloss were broken by the shuffled manipulation, which decreased perceived glossiness. Example images in **Figures 2B** (a and b) predominantly represent this aspect.

**Figure 2.**
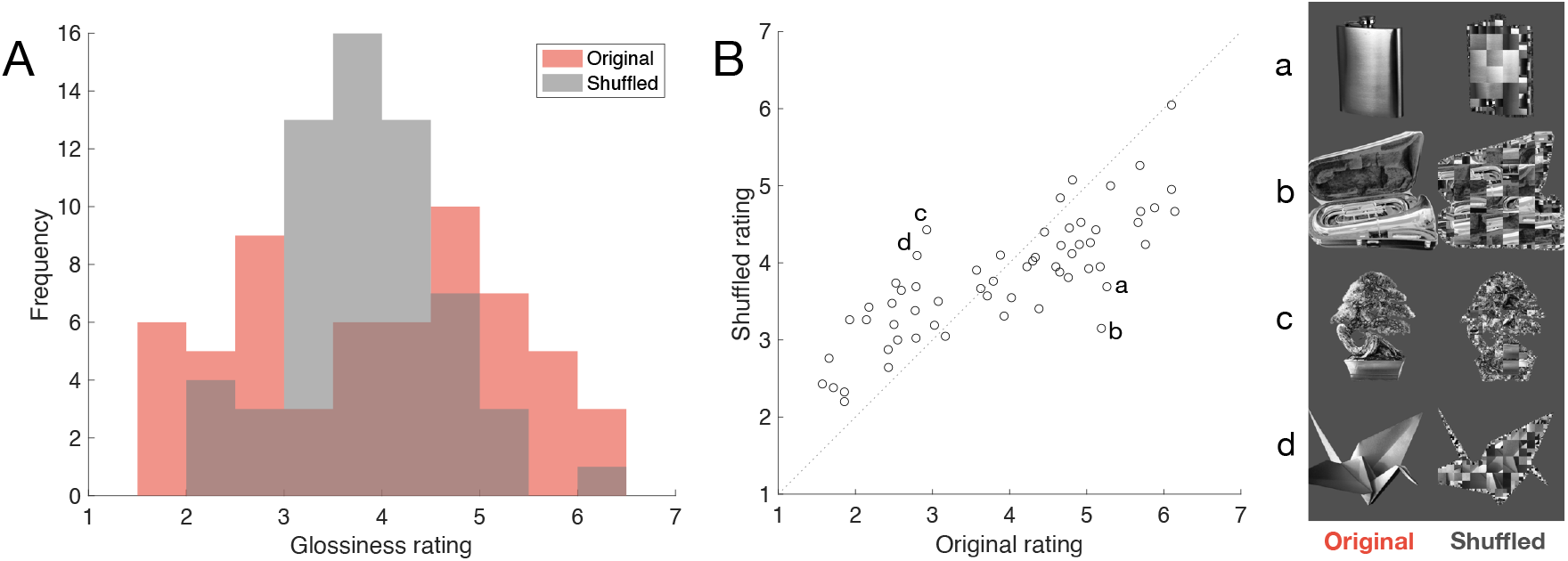
Glossiness rating. (A) Distribution of glossiness ratings of all stimuli. The horizontal axis indicates the average glossiness rating, and the vertical axis indicates the frequency with a 0.5 bin width. (B) Relationship between the original (horizontal) and shuffled (vertical) ratings. Each circle indicates each object image. Location on the diagonal dashed line indicates that the rating of the original and shuffled are the same. The right panel shows eight example images (a-d) with highly changed rating owing to the shuffling.

In contrast, we also found that shuffled manipulation caused high glossiness perception even when the original images were rated as low-glossiness objects (see example images in **Figure 2B** (c and d). The pixel arrangement of the images with low glossiness ratings did not accidentally constitute a structure imitating the specular highlights, but rather the magnitude of the perceived matte in the original object decreased by the shuffled manipulation. Therefore, these images were rated as glossy.

### Pupil diameter

**Figure 3A** shows the average pupillary responses for all original and shuffled images. Both indicate similar trends of pupil constriction when the stimulus onset is near 1,000 ms as the pupillary light reflex (PLR). Both peaks are nearly identical, indicating that the luminance of the stimuli was precisely controlled. However, the original and shuffled images thereafter form different curves. These findings imply that the original images were more constricted by some factors related to perceived glossiness or that the shuffled images were more dilated owing to some interest in the existing object. We propose that the gap between the original and shuffled was less affected by a difference in the physical intensities because their luminance histograms had been equalized. Instead, some cognitive factors could be driving these differences.

**Figure 3.**
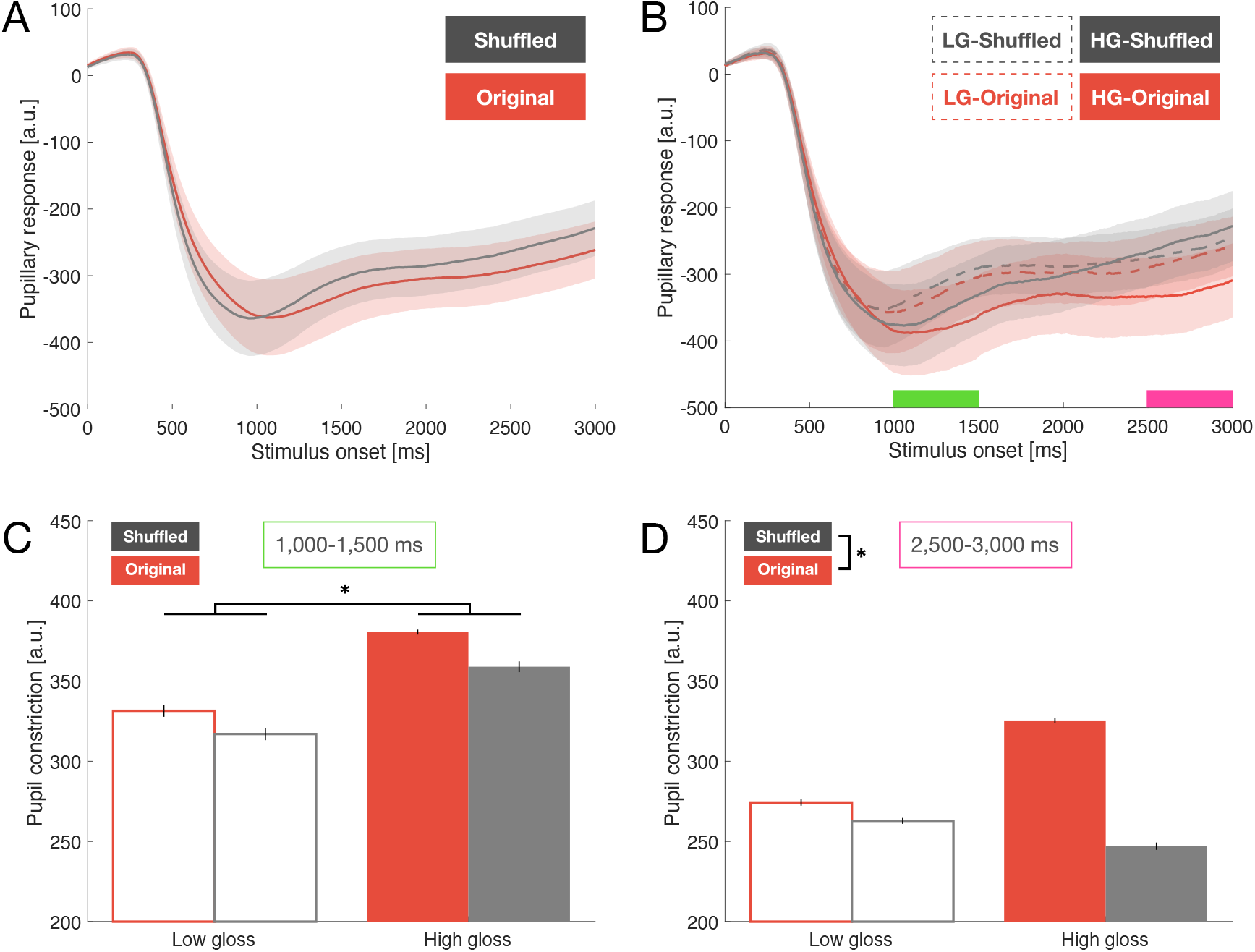
Pupillary responses. (A) Average pupillary responses for the original and shuffled conditions of all images (60 images in each). The horizontal axis indicates the stimulus onset, and the vertical axis indicates the pupillary response. Error bars represent the standard error of the mean. (B) Average pupillary responses grouped by the rating and image conditions (10 images in each). Green (1,000-1,500 ms) and pink (2,500-3,000 ms) areas illustrate intervals for further analysis shown in C and D, respectively. Other formats are the same as A. (C) Average pupil constriction in the early stage. The horizontal axis indicates four groups (LG-original, LG-shuffled, HG-original, HG-shuffled; from right to left). The vertical axis indicates the changes in pupil constriction, which is vertically opposite of B. (D) The same as C, except the averaging interval is in the later stage. Asterisks indicate statistically significant differences in the main effect of the ANOVA; * *p* < 0.05.

Therefore, we focused on the higher and lower glossiness images rated by the observers to directly discuss the relationship between perceived glossiness and pupillary responses. For each observer, the ten of each highest and lowest-rated original and shuffled images were extracted, which were defined as high gloss (HG) and low gloss (LG) images, respectively. **Figure 3B** shows that the HG-original and HG-shuffled were more constricted than the LGs at the PLR after approximately 1,000 ms from the stimulus onset. We presume that these different trends represent the dynamics of perceived glossiness (HG versus LG). In contrast, approximately 3,000 ms later, the HG- and LG-shuffled were more dilated than the originals, which suggests that these responses were derived from whether the object was recognized (original vs. shuffled).

Moreover, we computed the average pupillary diameters in each condition during two intervals. These were defined as early stage (1,000-1,500 ms), as show in **Figure 3C**, wherein the PLR strongly constricts (Mathôt, 2018), and later stage (2,500-3,000 ms), as shown in **Figure 3D**, wherein the pupil dilation reaches a peak affected by object recognition (Beukema et al., 2019). We performed two-way repeated measures analysis of variance (ANOVA) for the rating (HG vs. LG) and image conditions (original vs. shuffled) in the earlier stage. The main effect of the rating condition was significant (*F*(1, 20) = 6.13, *p* < 0.05, *n*_*p*_^2^ = 0.24), indicating that HG was more constricted than LG near the PLR (see **Figure 3C**). The image condition effect was insignificant (*F*(1, 20) = 1.24, *p* = 0.28, *n*_*p*_^2^ = 0.06), and there was no significant interaction (*F*(1, 20) = 0.10, *p* = 0.76, *n*_*p*_^2^ = 0.01). These results suggest that higher gloss images would constrain pupil size.

We similarly performed two-way repeated measures ANOVA in the later stage (see **Figure 3D**). The main effect of the image condition was significant (*F*(1, 20) = 7.40, *p* < 0.05, *n*_*p*_^2^ = 0.27), indicating that the shuffled was more dilated than the original. The rating condition effect was insignificant (*F*(1, 20) = 0.59, *p* = 0.45, *n*_*p*_^2^ = 0.03), and there was no significant interaction (*F*(1, 20) = 3.31, *p* = 0.08, *n*_*p*_^2^ = 0.14). Contrary to the early stage, the results in the later stage indicated a difference between the original and shuffled, which suggests that the shuffling operation affects pupil dilation.

## Discussion

This study investigated the relationship between glossiness perception and pupillary responses through a glossiness rating experiment that included pupil recordings. The stimuli included the original and shuffled images, wherein image features derived from the luminance histogram were controlled to the same values. We hypothesized that the pupillary responses were either constricted by cognitive factors related to higher glossiness perception or that there was no difference due to the control of the physical factors (the image features).

The experimental results indicate that when the original objects were rated as having high gloss, their shuffled images were rated lower (**Figure 2B**). This suggests that the shuffling process inhibited the global features of an image, such as the congruency of specular highlights, while retaining more local features. This finding supports previous studies showing that the congruency of specular highlights works as a cue for glossiness perception (e.g., Kim et al., 2011; P. Marlow et al., 2011; Todd et al., 2004). Conversely, when the original images were rated as having low gloss, their shuffled images were rated as having higher gloss. This inverse relation implies that we perceive “matteness” based on the global features broken by our pixel shuffled operation. It can be assumed that breaking global features not only decreases perceived glossiness but also increases perceived matteness, and such image editing would reveal a new verbalizable cue for matteness perception underlying the visual system.

Pupillary recording revealed that higher-gloss rated images were more constricted than lower-gloss rated images (**Figure 3C**). This suggests that not only the physical luminance but also the cognitive factors that the object surfaces were perceived as shinier constrict the pupil size, similar to subjective brightness (Binda et al., 2013b; Bombeke et al., 2016; Laeng et al., 2018; Laeng & Endestad, 2012; Naber & Nakayama, 2013). In other words, illusory brightness (Kinzuka et al., 2021; Suzuki, Minami, & Nakauchi, 2019; Suzuki, Minami, Laeng, et al., 2019; Zavagno et al., 2017) would appear in the images wherein glossiness perception primarily occurs. Specifically, for example, the visual system naturally attends brighter regions, eliciting pupil constriction (Binda et al., 2013a). A similar phenomenon occurs in the case of a surface’s specular highlight. With respect to pupil constriction, Tanaka et al. reported that the pupil size became smaller when the observers carefully observed the surface of the marble stone, even when the stimuli did not contain particularly brighter regions (Tanaka et al., 2017). Our results suggest that the specular reflection on the smooth surface of the stone stimulus can explain why the pupil was constricted in that case.

Even when the image features of the shuffled were the same as the original, the shuffled stimuli elicited more dilation than the original at a later stage (**Figure 3D**). This suggests that the observers may have had difficulty recognizing the object in the shuffled condition. Particularly, their pupils responded to unknown objects (Beukema et al., 2019) with some cognitive processing dilating the pupil diameter, such as mental workload (Klingner et al., 2011). Likewise, the shuffled images contained unfamiliar objects, leading to pupil dilation similar to the old/new effect (Kafkas & Montaldi, 2011; Otero et al., 2011). From a physiological perspective, one explanation for this reaction is that the locus colliculus (Nieuwenhuis et al., 2011) or intermediate layers of the superior colliculus (Wang et al., 2012, 2014) evoked these pupillary responses. Regarding arousal levels, Beukema and colleagues considered novelty and cognitive effort as possibilities for more dilation caused by unfamiliar objects (Beukema et al., 2019). It appears that the images of the shuffled condition were almost new, and it was difficult to identify the object. Thus, it is presumable that the observers’ pupillary responses in this study were caused by these processes.

Our results allow us to explain the dynamics of the difference between the two image attributes (the rating and image conditions). We observed that changes in pupil diameter based on perceived glossiness appeared near the PLR. This timing is similar to the top-down bias based on the shape from shading (Sapir et al., 2021). Thereafter, the changes in pupil diameter affected by image shuffling were observed at a later stage. This temporal alternation suggests that perceived glossiness is faster than the recognition of the object. Sharan et al. reported that material judgment is faster than real versus fake discrimination (Sharan et al., 2014), and our findings support this in the same temporal context. Similarly, Nagai et al. reported that identifying visual material features (e.g., glossiness and transparency) is faster than identifying non-visual material features (e.g., heaviness and hardness) in the material category discrimination task (Nagai et al., 2015). Here, the experience of touching or interacting with the object might play an important role in discrimination. Thus, in this study, such temporal features for material perception were verified from the aspect of pupillary responses. Furthermore, the continuous change in the cognitive factors from glossiness to object ambiguity perception implies the possibility of sharing networks with the pupil regulation system (Mathôt, 2018; Wang & Munoz, 2015) and the inferior temporal cortex related to gloss perception (Baba et al., 2021; Komatsu et al., 2021; Nishio et al., 2012, 2014). Therefore, further investigation through pupillometry is required to understand how glossiness perception and other cognitive factors are linked.

We controlled the image features based on the luminance histogram of all stimuli in this experiment. Its skewness was positive and the same value between the original and shuffled images (original: 1.24±0.08, shuffled: 1.27±0.09) was indicated, but they resulted in different perceived glossiness. Although positive skewness is a cue for high glossiness perception (Motoyoshi et al., 2007; Sharan, Li, et al., 2008), our results indicate that it contains both high and low glossiness perception. This suggests that more cognitive factors, such as memory related to the object, affect glossiness judgment in the visual system. It is possible that our pupillometry approach suitably captures the changes from these factors. An alternative aspect is that other cues, such as darker regions, contribute to perceived gloss (Kim et al., 2012; Kiyokawa et al., 2019).

We also consider whether any physical factors affected the pupillary response. Although the observers were instructed to focus on the fixation point at the center of the display, their line of sight may have been misaligned, and such a luminance difference would have affected the pupil diameter. Thus, we computed local luminance changes within one degree of the visual angle of the gaze point (Liao et al., 2020) at the early stage, based on recorded eye movements of the observers, and these changes and pupil size were not significantly correlated (*r* = 0.19, *p* = 0.24). Likewise, there was no significant correlation between the object area and pupil size (*r* = -0.19, *p* = 0.24). They suggested that such luminance changes on a pixel-by-pixel basis do not provide any meaningful differences as physical factors. Thus, these findings support our suggestion that the changes in pupillary response in the current study were mainly influenced by cognitive factors.

Although this experiment used general objects in the real world with low-or high-gloss surfaces, it is not clear how those objects’ semantic factors affected the gloss rating. For example, the image of a diamond (right column in **Figure 1A**) was rated as a high-gloss object that constricted the pupil size. It contains not only physically based surface properties, such as glossiness, but also more emotional or sensitive factors (e.g., preference, luxury, and desire). Such higher-level surface features are regarded as a kind of positive emotion and provide high-arousal stimulation, which leads to pupil dilation (Bradley et al., 2008), for example, perceived cuteness (Kuraguchi & Kanari, 2020, 2021). Thus, these stimuli would elicit both constriction and dilation owing to their physical and sensitive properties, respectively. Therefore, segregating a combination or hierarchy of these properties to further understand the visual process is highly recommended.

In conclusion, our findings suggest that the high-gloss rated images elicit constriction of the pupil size compared to the low-rated images, which is faster than the pupil dilation owing to object ambiguity. Further studies are required to explore the relationship between other perceptual qualities, such as transparency and the pupillary response, and whether the changes in the pupil responses depend on specific physical materials (e.g., wood, metal, glass). Moreover, connecting more emotional qualities beyond physical surface qualities with pupillometry would help us understand the dynamics of material perception in the visual system.

## Acknowledgements

This work was supported by JSPS KAKENHI, Grant Numbers JP21K21315, JP20H04273, and JP20H05956. The authors wish to thank Ryoya Shiomoto for supporting the data collection.

